# CRISPR-based editing reveals edge-specific effects in biological networks

**DOI:** 10.1101/265710

**Authors:** Yi Li, Chance M. Nowak, Daniel Withers, Alexander Pertsemlidis, Leonidas Bleris

**Affiliations:** Department of Bioengineering, The University of Texas at Dallas, 800 West Campbell Road, Richardson TX 75080 USA; Center for Systems Biology, The University of Texas at Dallas, 800 West Campbell Road, Richardson TX 75080 USA; Department of Biological Sciences, The University of Texas at Dallas, 800 West Campbell Road, Richardson TX 75080 USA; Greehey Children’s Cancer Research Institute, The University of Texas Health Science Center at San Antonio, 8403 Floyd Curl Drive, San Antonio TX 78229 USA

## Abstract

Unraveling the properties of biological networks is central to understanding normal and disease cellular phenotypes. Networks consist of functional elements (nodes) that form a variety of diverse connections (edges) with each node being a hub for multiple edges. Herein, in contrast to node-centric network perturbation and analysis approaches, we present a high-throughput CRISPR-based methodology for delineating the role of network edges. Ablation of network edges using a library targeting 93 miRNA target sites in 71 genes reveals numerous edges that control, with variable importance, cellular survival under stress. To compare the impact of removing nodes versus edges in a biological network, we dissect a specific p53-microRNA pathway. In summary, we demonstrate that network edges are critical to the function and stability of biological networks. Our results introduce a novel genetic screening opportunity via edge ablation and highlight a new dimension in biological network analysis.

## Introduction

We focus on the network formed by p53 and its upstream and downstream regulators, which is critical to cell health, yet incompletely understood. Since its discovery in 1979 (1), p53 has been shown to play a crucial role in maintaining genomic stability (2), with more than 50% of human cancers harboring mutant or deleted p53 (3). Under normal conditions, the p53 protein exists in a latent form and at low concentration, but in response to various cellular stress signals such as DNA damage, hypoxia, and oncogene expression, posttranslational modification of p53 results in its stabilization and accumulation (4). As most human malignancies shut down the p53 tumor-suppressing responses, p53 is one of the most promising targets for drug interventions in cancer therapy.

A class of post-transcriptional regulators, called microRNAs (miRNAs), is directly associated with p53, either regulating mRNA responsible for p53 production or being regulated by p53 and its partners (5, 6). miRNAs, in their mature forms, are small non-coding RNAs approximately 22 nucleotides in length that act as major regulators of gene expression. Since miRNAs are involved in critical cellular processes such as growth, differentiation, and apoptosis, the loss of critical miRNAs in a given cell type can have significant implications for cell fate (7–9). Studies have revealed extensive crosstalk between the p53 network and miRNAs, but the specifics of how miRNAs participate in the regulation of p53 signaling and what they contribute to the role of p53 as a tumor suppressor remain largely elusive.

In miRNA-based networks the edges are regulatory interactions between miRNAs and target mRNAs. These interactions are mediated by sequence complementarity and therefore are susceptible to genetic variation occurring in either the miRNA or the target site. Variation in miRNA binding sites has been associated with numerous diseases, including Tourette Syndrome (10), rheumatoid arthritis (11), lupus (12), psoriasis (13), Crohn’s disease (14, 15), Parkinson’s disease (16), hypertension (17, 18), diabetes and obesity (19, 20), and multiple cancers (21–25). In the context of p53 signaling (26), miR-34a regulates HDM4, a strong repressor of p53, creating a positive feedback loop in which high levels of miR-34a de-repress p53 which in turn transcriptionally up-regulates the expression of miR-34a.

Network edges (e.g. miRNA-gene target interactions) are therefore central to the function and stability of biological pathways, yet comprehensive studies focusing on edges remain few. Today, we have the unprecedented opportunity to dissect individual cells and pathways with single-nucleotide specificity using genome editing. The most widely-adopted editing methodology to date is the bacterial type II clustered regularly interspaced short palindromic repeats (CRISPR) system consisting of the CRISPR-associated protein Cas9 derived from *Streptococcus pyogenes* (SpCas9), a DNA endonuclease, and a guide RNA, which directs the binding of Cas9 to a DNA target upstream of a protospacer adjacent motif (PAM).

The CRISPR technology has revolutionized our ability to probe and edit the human genome *in vitro* and *in vivo,* including targeted gene disruption, insertion, single-nucleotide mutation, and chromosomal rearrangements (27–29). Furthermore, pooled sgRNA libraries can be used for versatile *in vitro* screening to investigate phenotypes of interest. Recent examples include screens identifying genes conferring drug resistance (30), genes involved in metastasis (31), and long non-coding RNAs (lncRNAs) regulating human cancer cell growth (32). Thus far, pooled sgRNA libraries have been applied to transcribed loci, which correspond to network nodes. Here, we selectively remove edges in the miRNA-p53 network, using a first-of-a-kind CRISPR-based screen. We demonstrate that removing edges sheds new light on pathways, in ways not achievable through node-based approaches, which may lead to novel and non-obvious therapeutic opportunities.

## Results

### CRISPR-based screen for microRNA target editing

We focus on five miRNAs – miR-34a, miR-145, miR-192, miR-194 and miR-215 – which are known to be directly or indirectly regulated by p53, and play elaborate roles in the p53 pathway ((33, 34), review (35)). The target genes for each miRNA were compiled from miRTarBase (36). We selected targets that have been experimentally validated by multiple methods, including luciferase reporter assay, western blot, and quantitative RT-PCR (qRT-PCR) (**Supplementary Table 1**). For each of the target genes, the miRNA target sites within its 3’UTR were determined using TargetScan (37). In total, 93 miRNA target sites were identified across the 71 target genes. The miRNAs and the 71 target genes are the nodes of the derived network, while the experimentally verified and high-confidence predicted interactions between the nodes, including interactions between miRNAs and target genes and between target genes themselves, are the network edges (Figure 1a). Since SpCas9-mediated NHEJ typically introduces short insertions or deletions (indels) near its cutting site, we designed sgRNAs in which a PAM is adjacent to the miRNA target seed sequences within the 3’UTR (38) **(Supplementary Table 1)**.

**Figure 1.**
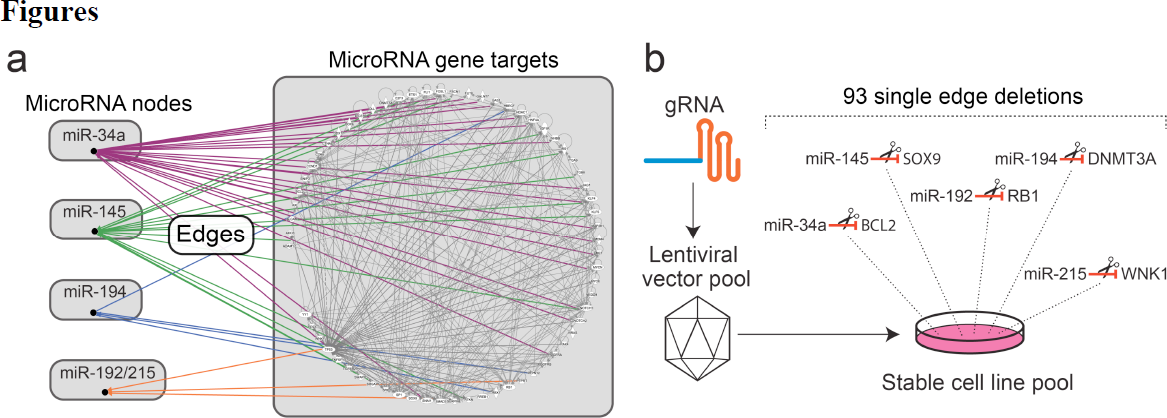
p53-miRNA network and CRISPR-based edge screens. **(a)** Complexity of the p53-miRNA network with nodes comprising the indicated miRNAs and their 71 target genes and edges based on experimentally verified and high-confidence predicted direct interactions, derived using Qiagen Ingenuity Pathway Analysis. **(b)** CRISPR-based lentiviral libraries were prepared using the lentiCRISPRv2 system. The stably integrated CRISPR sgRNA constructs were recovered by PCR and the sgRNA targets were identified using NGS.

Next, we constructed a pooled CRISPR sgRNA library, containing both SpCas9 and sgRNA expression cassettes (39). Equimolar amounts of the 93 pairs of sgRNA oligonucleotides were mixed and cloned into a lentiviral vector (lentiCRISPRv2). The resulting plasmid library was subjected to sequencing, to confirm library complexity. The resulting reads displayed a 20-bp “noisy” sgRNA target sequence matching the expected pattern from the sgRNA mixture **(Supplementary Figure 1)**. Subsequently, the lentiviral library was used to infect HCT116 wild-type (WT) and HCT116 p53^-/-^ cells at a multiplicity of infection (MOI) of 0.3, which has been shown (40) to yield at most one integration of the sgRNA cassette in the majority of cells (**Figure 1b**). To verify the complexity of our resulting libraries in cells (named LIB-WT and LIB-p53"^-/-^") was maintained, the sgRNA locations were amplified from genomic DNA and subjected to Sanger sequencing, which again displayed the expected pattern **(Supplementary Figure 2)**.

In parallel, to test the efficacy of the viral system, we prepared two CRISPR lentiviral vectors that target the open reading frame (ORF) of the zeocin resistance gene (target 1: 5’-TCGCCGGAGCGGTCGAGTTC-<u>TGG</u>; target 2: 5’-CTCACCGCGCGCGACGTCGC-<u>CGG</u>; PAM underlined), and stably integrated them into cell line Flp-In-293 which harbors the zeocin resistance gene (ThemoFisher Scientific). As shown in **Supplementary Figure 3**, disruption of the Zeocin resistance gene abolished resistance to Zeocin (100 μg/mL) in the two resulting cell lines (FLP-EDIT1 and FLP-EDIT2), compared to the parental Flp-In-293 cells.

Using the established cell lines (LIB-WT and LIB-p53^-/-^), we focused on the role of miR-34a in the overall p53-miRNA network (**Figure 1a**). miR-34a is transcriptionally activated by p53 and induces an antiproliferative phenotype including senescence, cell cycle arrest at the G1 phase, and apoptosis (41, 42). In turn, overexpression of miR-34a increases p53 protein level and stability (33). Importantly, our established cell lines (LIB-WT and LIB-p53^-/-^) have low-level baseline miR-34a expression **(Supplementary Figure 5)** while the ectopic miR-34a delivery results, on average, in a 71-fold increase.

We adopted a survival assay mediated by ectopic exposure to miR-34a mimics. Specifically, we treated both cell lines (LIB-WT and LIB-p53^-/-^) to 25 nM of miR-34a mimic for 6 days. Cells were harvested at day 0 (before miRNA treatment) and at day 6. For each sample, sgRNA constructs were amplified from genomic DNA and subjected to the Illumina NGS Amplicon Sequencing to assess the relative abundance for each of the 93 sgRNA target sequences **(Supplementary Table 2)**. The most enriched or depleted sgRNA targets, defined by fold-changes between Day 6 and Day 0 larger than 10, were identified for both LIB-WT and LIB-p53^-/-^ cells **(Figure 2, Supplementary Table 3)**.

**Figure 2.**
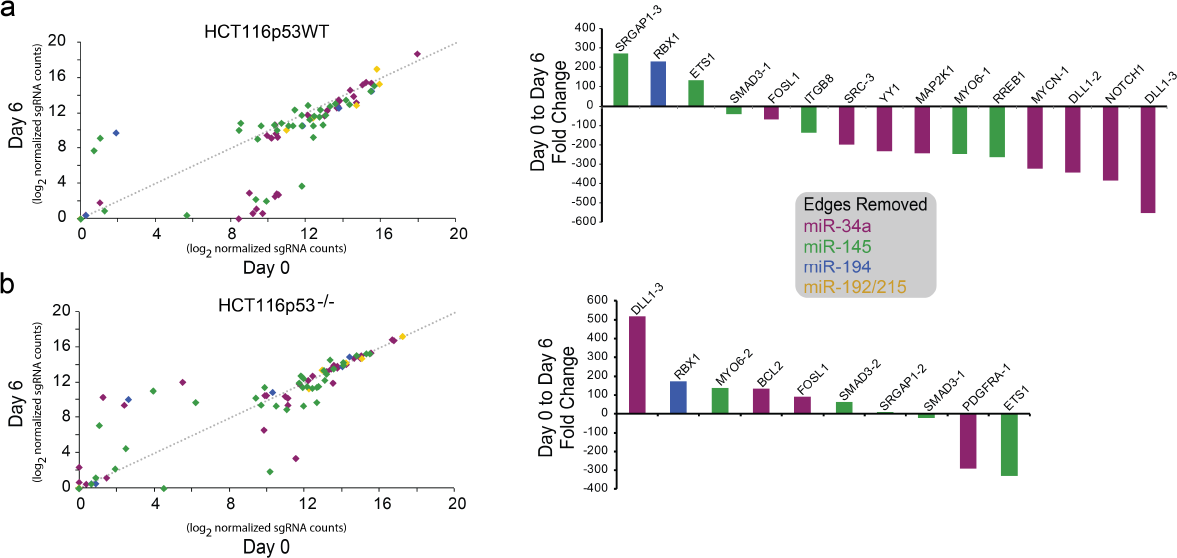
High-throughput editing of edges with CRISPR libraries in (a) HCT116p53WT (LIB-WT) cells and (b) HCT116p53^-/-^ (LIB-p53^-/-^). The sgRNA targets showing the highest folder changes (> 10) after 6 days treatment of 25nM with miR-34a mimic are shown, with positive values indicating enrichment and negative values indicating depletion.

Intriguingly, RBX1 (RING-box protein 1), a RING subunit of SCF (Skp1, Cullins, F-box) E3 ubiquitin ligases, was highly enriched in both cell lines. Although not a direct target of miR-34a, overexpression of RBX1 has been demonstrated to increase cancer cell survival (43), and thus could serve as a general response mechanism to cellular stress induced by ectopic miR-34a. Additionally, for a subset of gene targets, we observed differential response to miR-34a treatment between the LIB-WT and LIB-p53^-/-^ cells **(Supplementary Table 3)**. For example, the sgRNA targeting the anti-apoptotic gene Bcl-2 was enriched in the LIB-p53^-/-^ cells after miR-34a treatment while no enrichment was observed in the LIB-WT cells **(Supplementary Figure 4)**.

### Node perturbations versus edge edits in biological networks

Our edge editing approach revealed **(Figure 2b)** several clones that are enriched or depleted after prolonged exposure to ectopic miR-34a. To assess the impact of edge removal (through ablation of miRNA:target interactions) we focused on Bcl-2, a gene that in response to miR-34a has different response between the two cell lines **(Figure 2b)** and is known to be involved in cell survival (44).

Returning to the HCT116 wild-type (WT) and HCT116 p53^-/-^ cells, we removed the miR-34a target site from the Bcl-2 locus. We prepared a single sgRNA construct designed against the Bcl2 3’UTR and established stable cell lines (BCL2tgt-WT and BCL2tgt-p53^-/-^) using the same viral delivery system. Sanger sequencing of PCR products spanning the sgRNA target site showed that edits (indels) occurred immediately upstream of the PAM **(Supplementary Figure 6**, AGG, highlighted) in both cell lines.

Treating delivery of ectopic miR-34a mimic as perturbation of a network node and removal of the miR-34a/Bcl-2 interaction as perturbation of a network edge **(Figure 3a)**, there are four possible combinations (node and edge present/absent). When mir-34a levels are low (i.e. the node is absent) the presence or absence of the edge does not impact survival **(Supplementary Figure 7)** of either cell line.

**Figure 3.**
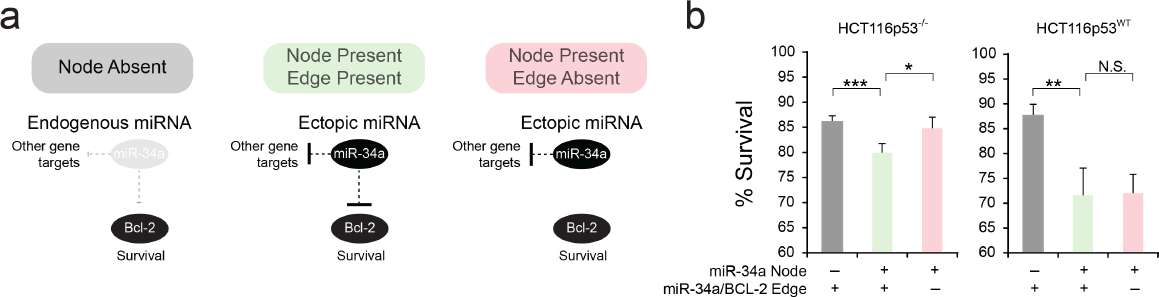
Survival analysis after ectopic miR-34a delivery using node vs. edge approaches. (a) Schematic illustration of node vs. edge analysis. Ectopic miR-34a represents a network node and the miR-34a/Bcl-2 interaction represents a network edge. (b) The node-based approach shows that addition of the miR-34a node induces apoptosis in both p53 WT and deficient cells. In contrast, the edge-based approach reveals that introduction of the miR-34a/Bcl-2 edge induces apoptosis only in the p53-deficient cells and not in p53-WT cells.

In the context of node perturbation, the introduction of ectopic miR-34a in wild-type cells induces apoptosis **(Figure 3b right panel and Supplementary Figure 8**; cell viability is 87.6% without the miR-34a node, and 71.6% with the miR-34a node, with a p-value of 0.006). Similar changes were observed in p53^-/-^ cells (**Figure 3b left panel** and **Supplementary Figure 8**; cell viability is 86.1% without the miR-34a node, and 80.0% with the miR-34a node, with a p-value of 0.002). In this case, perturbing the miR-34a node results in the same behavior for both p53 wild-type and p53^-/-^ cells.

In the context of edge perturbation, we observe that the response of the two cell lines to ectopic miR-34a is sensitive to the presence of the miR-34a/Bcl-2 edge. Specifically, removing the ability of miR-34a to regulate Bcl-2 in the p53^-/-^ cells induces apoptosis **(Figure 3b left panel and Supplementary Figure 8**; cell viability is 84.8% without the miR-34a/Bcl-2 edge, and 80.0% with the miR-34a/Bcl-2 edge, with a p-value of 0.015), while no such phenotypic changes are observed in p53 WT cells **(Figure 3b right panel and Supplementary Figure 8**; cell viability is 72.0% without the miR-34a/Bcl-2 edge, and 71.6% with the miR-34a/Bcl-2 edge, with a p-value of 0.90). We note that the same conclusions can be drawn when quantifying the early-or late-apoptotic cells **(Supplementary Figure 9)**.

To further explore the response to miR-34a in BCL2tgt-WT cells, we quantified expression of the Bcl-2 mRNA in response to miR-34a treatment using qRT-PCR. As expected, miR-34a suppresses the expression of Bcl-2 mRNA in the WT and p53^-/-^ cells by 55% and 40%, respectively **(Supplementary Figure 10)**. In the BCL2tgt-p53^-/-^ cells, ectopic miR-34a has a minimal effect on Bcl-2 mRNA level (95% compared to the control-treated sample, with a p-value of 0.71). In the BCL2tgt-WT cells, ectopic miR-34a results in a significant down-regulation of Bcl-2 expression (62% compared to the control-treated sample, with a p-value of 0.028), possibly due to additional p53-miR-34a regulatory mechanisms.

## Discussion

Biological networks consist of nodes and the interactions between them (i.e. edges). Conventional screening methods remove one node at a time, disrupting all edges connected to that node, and therefore producing a relatively blunt effect. An inhibitor that perturbs or removes a single node yields diverse and systemic changes in the whole network through both direct and indirect connections (45), which may explain the heterogeneity observed with single-molecule associated therapeutics.

Our approach reveals edge-specific effects related to the pro-apoptotic p53 and anti-apoptotic Bcl-2 proteins, focal nodes of apoptotic signaling. Normally, p53-dependent inhibition of Bcl-2 and induction of BAX, PUMA and NOXA overcome the anti-apoptotic threshold set by Bcl-2 family members. Conceivably, the difference in apoptosis observed in the BCL2tgt-WT and BCL2tgt-p53^-/-^ cells under miR-34a treatment **(Figure 3b)** may be explained by the presence of wild type p53-dependent upregulation of PUMA or NOXA in p53 WT cells and not in p53^-/-^ cells. Additionally, p53 could disrupt the binding of POU4F1 to the promoter of Bcl-2 and thus indirectly down-regulate Bcl-2 expression **(Supplementary Figure 10)**.

Taken together, our results show that the disruption of the edge between miR-34a and Bcl-2 can revert the miR-34a triggered apoptotic effects in a p53-deficient cell model. In WT cells, introducing miR-34a presumably indirectly triggers the suppression of the expression of Bcl-2 via an alternative p53-related pathway, which subsequently leads to increased apoptosis.

In conclusion, our CRISPR-mediated edge screening can be used to dissect critical biological interactions essential to cell survival. More generally, we demonstrate that the subtle effect of our edge removal methodology offers superior resolution and granularity in the analysis of biological networks and can lead to the identification of previously hidden interactions and opportunities for intervention.

## Materials and Methods

### Preparation of the CRISPR plasmid library

The CRISPR plasmid library was prepared by following the lentiCRISPRv2 cloning protocol provided by Dr. Feng Zhang (Department of Biology, MIT). Briefly, for each identified sgRNA target (20 nt), two oligos were synthesized. The first oligo was designed as 5’-CACCG-(20 nt sgRNA target sequence)-3’. The second oligo was designed as 5’-AAAC-(20 nt reverse complement of the sgRNA target sequence)-C-3’. All 93 pairs of oligonucleotides were synthesized by Sigma-Aldrich using its customized 96-well plate format **(Supplementary Table 4)**. Each well contained the pair of oligos for a specific sgRNA target (100 nmol for each). The oligo pairs were reconstituted using 100 μL of dH_2_O. For annealing the oligo pairs, 2 μL of each of the reconstituted oligo solutions was mixed with 2 μL of 10X T4 DNA Ligase Buffer (New England Biolabs, catalog number: B0202S) and 16 μL dH_2_O. The mixtures were heated at 95°C for 4 minutes, then left at room temperature for 60 minutes. 1 μg of the lentiCRISPRv2 plasmid (Addgene, catalog number: 52961) was digested with 1 μL Esp3I (ThermoFisher Scientific, catalog number: ER0451) at 37°C for 1 hour and run out on an X% agarose gel. The 12 kb band was extracted using QIAquick Gel Extraction Kit (Qiagen, catalog number: 28704). 1 μL of each of the annealed oligo pairs was mixed with 9,904 μL dH_2_O. Subsequently, 1 μL of the oligo mixture was ligated with Esp3I-digested lentiCRISPRv2 using T4 DNA Ligase (New England Biolabs, catalog number: M0202S). To ensure the integrity of library complexity, XL10-Gold Ultracompetent cells (Agilent, catalog number: 200314) were transformed, and more than 300 individual clones were pooled together to prepare the CRISPR plasmid library. To confirm library complexity, the plasmid library was subjected to Sanger sequencing (Genewiz) using primer P1 and analyzed using FinchTV (Geospiza).

### Generation of the CRISPR lentiviral screen library

To generate the lentiviral vectors, HEK293T cells were grown to 50–70% confluence and then transfected with 3.3 pg of the CRISPR plasmid library, 3.3 μg of the pMD2-VSVG plasmid, and 3.3 μg of the psPAX2 plasmid using 20 mL of JetPRIME (Polyplus, catalog number: 114-01). 24 h later, the medium was removed and replenished with 5 mL of complete growth medium. In the next 3 days, the growth medium containing lentiviral vectors was harvested, and 5 mL of fresh complete growth medium was replenished. The final pooled 15 mL growth medium was centrifuged at 3,000 rpm for 15 min at 4°C to remove cell debris. The supernatant was filtered through a 0.45 μm filter, dispensed into 1–2 mL aliquots and stored at -80°C. Viral titers were determined using qPCR Lentivirus Titration Kit (ABMGood, catalog number: LV900) following manufacturer’s instructions. Briefly, 2 μL of viral stock was mixed with 18 μL of Virus Lysis Buffer and incubated at room temperature for 3 minutes. This viral lysate, together with positive control (STD1), positive control (STD2), and negative control (NTC), were subjected to qRT-PCR. Finally, the titer of the viral stock was calculated based on the formula provided by the manufacturer and determined to be 2.07× 10^7^ IU/mL. To generate the LIB-WT and LIB-p53^-/-^ stable cells, ~10 million cells were seeded onto a 10 cm petri dish. 16 hours later, cells were transduced using the lentiviral vectors at a multiplicity of infection (MOI) of 0.3. 48 hours post-transduction, cells were treated with 0.5 μg/mL of puromycin (ThermoFisher Scientific, catalog number: A1113802). Polyclonal stable cell line libraries were established after ~2 weeks of drug selection.

### Sanger amplicon sequencing

To confirm the complexity of the LIB-WT and LIB-p53^-/-^ cell line libraries, total genomic DNA was isolated from LIB-WT and LIB-p53^-/-^ cells using the DNeasy Blood & Tissue Kit (Qiagen, catalog number: 69504). The cDNA fragments harboring the sgRNA target sequences were PCR amplified by using ~100 ng of the genomic DNA and primers P2 and P3. PCR conditions were one cycle of 30 seconds at 98°C, 40 cycles of 10 seconds at 98°C, 30 seconds at 60°C, and 30 seconds at 72°C. The 181 bp product was then subjected to direct Sanger sequencing using primer P2 and analyzed using FinchTV (Geospiza). To determine editing efficiency, total genomic DNA was isolated from BCL2tgt-WT and BCL2tgt-p53^-/-^ cells using the DNeasy Blood & Tissue Kit (Qiagen, catalog number: 69504). cDNA fragments harboring the miR-34a target site within the 3’UTR of BCL2 were PCR amplified by using ~100 ng of genomic DNA and primers P8 and P9. The 191 bp product was then subjected to direct Sanger sequencing using primer P9 and analyzed using FinchTV (Geospiza).

### Next generation sequencing

To determine the relative abundance of the 93 sgRNA target sequences before and after the CRISPR screen, total genomic DNA was isolated from miR-34a-treated LIB-WT and LIB-p53^-/-^ cells at days 0 and 6 using the DNeasy Blood & Tissue Kit (Qiagen, catalog number: 69504). cDNA fragments harboring the sgRNA target sequences were PCR amplified by using ~100 ng of the genomic DNA and primers P10 and P11, which added the 5’-overhang adapter sequence (5’-TCGTCGGCAGCGTCAGATGTGTATAAGAGACAG-3’) and the 3’- overhang adapter sequence (5’-GTCTCGTGGGCTCGGAGATGTGTATAAGAGACAG-3’) for subsequent Illumina NGS amplicon sequencing, which was performed at the Genome Sequencing Facility (GSF) at The University of Texas Health Science Center at San Antonio (UTHSCSA). 2 million individual reads were generated for each sample. Subsequently, the relative abundances of all 93 sgRNA target sequences were calculated and represented as counts per million reads (CPM). Log-transformed values were used for presentation.

### Apoptosis assay

To determine the non-apoptotic cell population 72 hours post-treatment with miR-34a mimic, 1 mL of the original cell growth medium was transferred into a 15 mL conical tube. Cells were washed with 1 mL of PBS solution, which was also collected. Next, cells were trypsinized using 150 μL of trypsin-EDTA for 5 minutes at 37°C. Subsequently, the trypsin-EDTA was neutralized using 2 mL of the original cell growth medium/PBS washing solution mixture. The cells were harvested by centrifugation at 1,000 rpm for 5 minutes. The cell pellet was then re-suspended in 1 mL PBS solution, then subjected to centrifugation at 1,000 rpm for 5 minutes. Apoptosis was quantified using the Dead Cell Apoptosis Kit with Annexin V Alexa Fluor™ 488 & Propidium Iodide (PI) (Invitrogen, catalog # V13241), following manufacturer’s instructions. Briefly, the harvested cell pellets were re-suspended in 100 μL of 1X annexin-binding buffer before being stained with 1 μL propidium iodide (100 μg/μL) and 5 μL of stock annexin V, Alexa Fluor™ 488 conjugate for 15 minutes in the dark. Stained cells were then diluted with 400 μL of 1X annexin-binding buffer before subjected to flow cytometry. Excitation/emission wavelengths for the annexin V, Alexa Fluor™ 488 conjugate are 495/519 nm; for propidium iodide they are 533/617 nm.

### Cell viability assay

Approximately 150,000 of the HEK293, Flp-In-293, FLP-EDIT1 and FLP-EDIT2 cells were seeded into 6-well plates in 2 mL of complete medium. FLP-EDIT1 and FLP-EDIT2 cells were maintained with 0.5 pg/mL puromycin. All cells were treated with 100 μg/mL zeocin. For each cell type, 6 wells were included so that one of the wells could be harvested and counted on each day (from day 1 to day 6 after seeding). Three independent experiments were performed. For live cell counting, the cell suspension was mixed with 0.4% Trypan Blue solution (Invitrogen, catalog number: 15250) at a 1:1 ratio (volume:volume). Unstained, live cells were then counted using a hemocytometer (Hausser Scientific, catalog number: UX-79001-00) under a light microscope.

### Quantitative reverse transcription-PCR (qRT-PCR)

For measurement of BCL2 mRNA levels, total RNA was extracted using the RNeasy Mini Kit (Qiagen, #74104) 48 hours post-transfection. First strand synthesis was performed using the QuantiTect Reverse Transcription Kit (Qiagen, #205311). Quantitative PCR was performed using the KAPA SYBR FAST Universal qPCR Kit (KAPABiosystems, #KK4601), with GAPDH levels used for normalization. Quantitative analysis was performed using the 2^-ΔΔCt^ method. Fold-change values are reported as mean with standard deviation. Primers used for BCL2 were (P4) 5'- CATGCTGGGGCCGTACAG-3′ and (P5) 5′-GAACCGGCACCTGCACAC-3′. Primers used for GAPDH were (P6) 5'- AATCCCATCACCATCTTCCA-3′ and (P7) 5′-TGGACTCCACGACGTACTCA-3’.

### General cloning protocol

Q5 High-Fidelity 2X Master Mix (New England Biolabs, catalog number: M0492) was used for all polymerase chain reactions (PCR) according to the manufacturer’s protocol. All oligonucleotides were ordered from Sigma-Aldrich and are listed in Supplementary Table 5. Plasmids were constructed using PCR amplification, restriction digest (all restriction enzymes were ordered from New England Biolabs), and ligation with T4 DNA ligase (New England Biolabs, catalog number: M0202S). Gel purification and PCR purification were performed with QIAquick Gel Extraction (catalog number: 28704) and PCR Purification kits (catalog number: 28104) (Qiagen). Transformations were performed using NEB 5-alpha electrocompetent *Escherichia Coli* (New England Biolabs, catalog number: C2987P). Minipreps were performed using QIAprep Spin Miniprep kit (Qiagen, catalog number: 27104). The final plasmids were confirmed by both restriction enzyme digestion and direct Sanger sequencing.

### DNA Constructs

#### U6-BCL2/sgRNA-PEF1

A pair of oligonucleotides (Forward: 5’-CACCGAATCAGCTATTTACTGCCAA-3’, Reverse: 5’-AAACTTGGCAGTAAATAGCTGATTC-3’) were annealed and cloned into the Esp3I-treated lentiCRISPRv2 plasmid. For more details, refer to Methods: Preparation of the CRISPR plasmid library.

#### U6-zeocin resistance gene/sgRNA1-PEF1 and U6-zeocin resistance gene/sgRNA2-PEF1

The zeocin resistance gene/sgRNA1 and zeocin resistance gene/sgRNA2 sequences were prepared from two rounds of PCR: first, the fragments were PCR amplified from U6-BCL2/sgRNA-PEF1 using primers P12 and P13 or P14; second, the final fragments were PCR amplified from the first round PCR products using primers P12 and P15. Subsequently, the PCR products were cloned into the U6-BCL2/sgRNA-PEF1 plasmid using Kpnl and EcoRI sites.

### Cell culture and transient transfection

HEK293 cells were acquired from American Type Culture Collection (ATCC, catalog number: CRL-1573). HCT116 wild-type and HCT 116 p53^-/-^ cells were gifts from Dr. Michael A. White (University of Texas Southwestern Medical Center). Flp-In-293 cells were purchased from ThermoFisher Scientific (catalog number: R75007). All cell lines were maintained at 37°C, 100% humidity and 5% CO_2_. Cells were grown in Dulbecco’s modified Eagle’s medium (DMEM, Invitrogen, catalog number: 11965-1181) supplemented with 10% Fetal Bovine Serum (FBS, Invitrogen, catalog number: 26140), 0.1 mM MEM non-essential amino acids (Invitrogen, catalog number: 11140-050), and 0.045 units/mL of Penicillin and 0.045 units/mL of streptomycin (Penicillin-Streptomycin liquid, Invitrogen, catalog number: 15140). In addition, 100 μg/mL zeocin (ThermoFisher Scientific, catalog number: R25001) was used for maintaining the Flp-In-293 cells. To pass the cells, adherent cultures were first washed with PBS (Dulbecco’s Phosphate Buffered Saline, Mediatech, catalog number: 21- 030-CM), then trypsinized with Trypsin-EDTA (0.25% Trypsin with EDTAX4Na, Invitrogen, catalog number: 25200) and finally diluted in fresh medium. For transient transfection of miRNA mimics, 200 μL of DMEM was mixed with 25 nM (final concentration) of miR-34a mimic (Qiagen, catalog number: MSY0000255) or miR-cel-67 mimic (Dharmacon, catalog number: CN-001000-01), in addition to 2 μL of RNAiMAX (Invitrogen, catalog number: 13778030). The mixture was added to each well of 12-well culture treated plastic plates (Greiner Bio-One, catalog number: 665180) and incubated at room temperature for 20 minutes. Adherent cell cultures were washed with PBS, then trypsinized with Trypsin-EDTA and finally diluted in fresh medium to the cell density of 200,000 cells/800 μL medium. 800 μL of the diluted cell suspension was then added to the well containing the miRNA-RNAiMAX complex.

## Acknowledgements

This work was funded by the US National Science Foundation (NSF) CAREER grant 1351354, NSF 1361355, and the University of Texas at Dallas. We thank S. Lawson for technical support.

## Author contributions

Y.L. and L.B. designed the experiments. Y.L., C.N., and D.W. performed the experiments. Y.L., C.N., A.P, and L. B. analyzed the data. Y.L., C.N. A.P, and L.B. wrote the paper. L.B. supervised the project.

## Additional information

Supplementary information is available in the online version of the paper. Correspondence and requests for materials should be addressed to L.B.

